# Role of molecular polymorphism in defining tau filament structures in neurodegenerative diseases

**DOI:** 10.1101/2021.05.24.445353

**Authors:** Xinyu Xiang, Tamta Arakhamia, Yari Carlomagno, Shikhar Dhingra, Manon Thierry, Michael DeTure, Casey N. Cook, Dennis W. Dickson, Leonard Petrucelli, Anthony W. P. Fitzpatrick

## Abstract

Misfolding and aggregation of tau protein is implicated in many neurodegenerative diseases that are typified by the presence of large, filamentous tau inclusions. The aggregation of tau in human brain is disease-specific with characteristic filaments defining the neuropathology. An understanding of how identical tau isoforms aggregate into disparate filament morphologies in phenotypically distinct tau-related diseases remains elusive. Here, we determine the structure of a brain-derived twisted tau filament in progressive supranuclear palsy and compare it to a dissimilar tau fold found in corticobasal degeneration. While the tau filament core in both diseases is comprised of residues 274 to 380, molecular-level polymorphism exists. Potential origins of the molecular polymorphism, such as noncovalent cofactor binding, are identified and predicted to modulate tau filament structures in the brain.

---

Neurodegenerative diseases are characterized by the deposition of filamentous protein aggregates in the brain (*1*). Misfolded proteins, such as the microtubule-associated phosphoprotein tau, accrue intracellularly into tangled inclusions manifesting in numerous diseases known collectively as tauopathies (*2*). As a result of alternative splicing, six isoforms of tau are expressed in the human brain, broadly classified on the constituent number of microtubule binding repeats (3- or 4-repeats) and distinguished by the length of the N-terminal region (*3, 4*). It has been demonstrated that isoform composition largely determines tau filament structure in Alzheimer’s disease (AD, 3- and 4-repeats) (*5*), Pick’s disease (3-repeat) (*6*), and corticobasal degeneration (CBD, 4-repeat) (*7, 8*) and mediated by posttranslational modifications (*7*), thus preserving specificity in templated-filament growth, or seeding (*9*). However, it is not known how filaments formed by the same tau isoforms differ structurally and how this arises. Here, we solve the structure of a filament formed by tau in progressive supranuclear palsy (PSP), a tauopathy similar to CBD in which only 4-repeat (4R) isoforms accumulate and aggregate (*10, 11*). We find that 4R tau misfolds into two structurally distinct molecular shapes, or conformers, in PSP and CBD and discuss the possible origins of this molecular-level polymorphism (*12*).

In PSP brain (Fig. 1A), tau filamentous inclusions constitute neurofibrillary tangles and neuropil threads, associated in variable proportions with tufted and thorny astrocytes, as well as with oligodendroglial coiled bodies (Fig. 1, B and C) (*13, 14*). The burden of tau lesions attributed to PSP, shown by tissue staining (Fig. 1, B and C) and immunolabeling (Fig. 1D), is correlated with clinical symptoms (*15*). A tau filament morphology identified in many PSP cases is described as a twisted ribbon with a helical pitch of ~1,050 Å (Fig. 1E), a maximum width of ~150 Å, and a minimum width of ~70 Å (Fig. 1F) (*16*). Further, it has been determined that the highly stable, trypsin-resistant filamentous core is composed of tau fragments comprised of residues ~268-395, similar to CBD (*17*). We have determined the density map of a twisted tau filament morphology characteristic of PSP using cryo-electron microscopy (cryo-EM) to an overall resolution of 3.9 Å (Fig. 1G and fig. S1; see Methods for details).

**Fig. 1.**
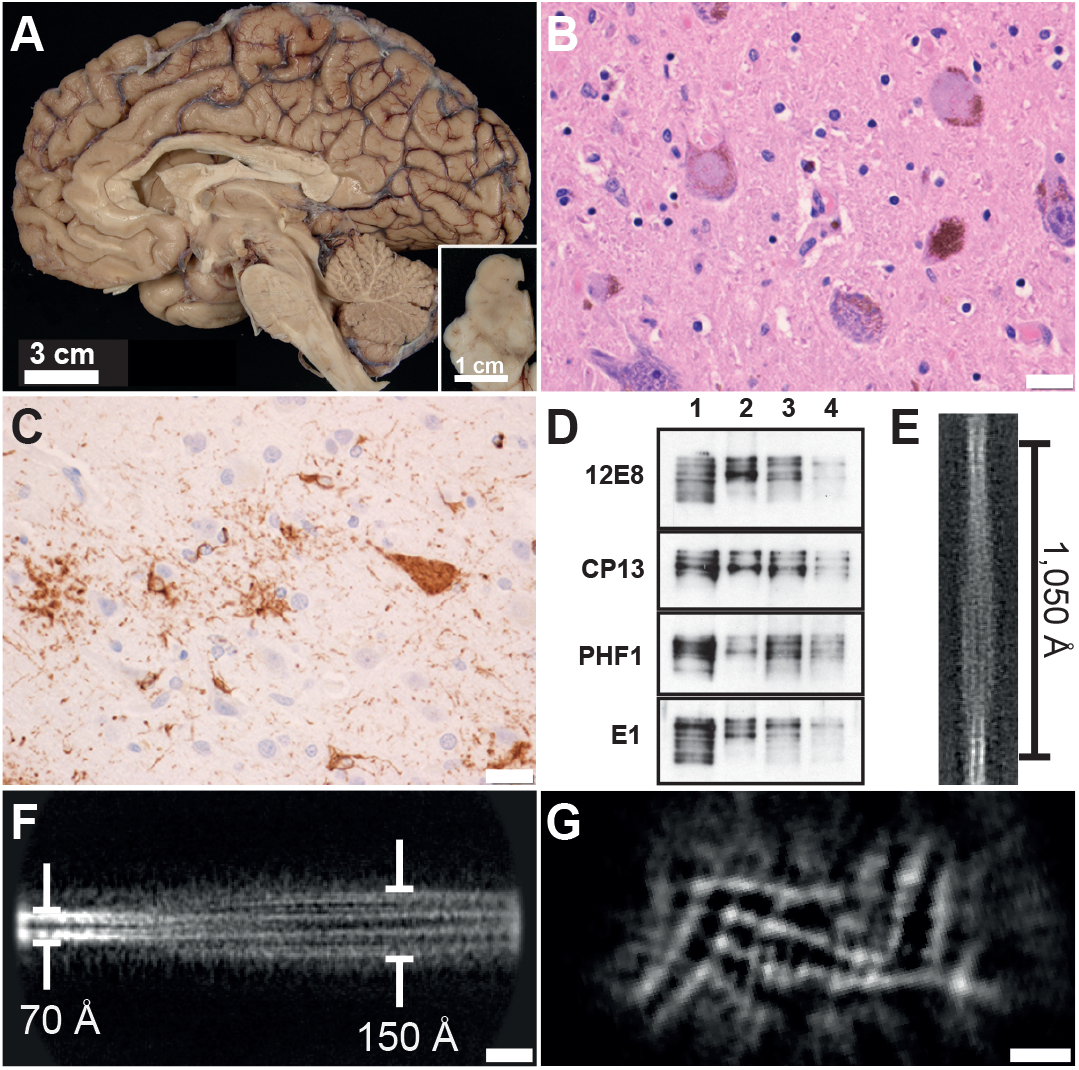
Neuropathological, biochemical, and cryo-EM analysis of twisted PSP tau filaments. A) Gross appearance of the PSP brain (mid-sagittal) shows midbrain atrophy and loss of neuromelanin pigment in the substantia nigra (inset). B) and C) The pigment-containing neurons of the substantia nigra have globus neurofibrillary tangles (hematoxylin and eosin), and tau immunohistochemistry shows neuronal and glial lesions (pretangles, tufted astrocytes, and oligodendroglial coiled bodies). Scale bar, 20 µm. D) Sarkosyl-insoluble fraction extracted from the caudate/putamen of four different PSP cases as evaluated by immunoblotting for the tau antibodies 12E8 (pS262/356), CP13 (pS202), PHF1 (pS396/404), and E1 (human tau-specific). Tau filaments from PSP case #1 were subsequently used for cryo-EM analysis. E) 2D class average of PSP tau filaments at 5.3025 Å/pixel with a helical pitch of ~1,050 Å. F) 2D class average of PSP tau filaments at 3.1815 Å/pixel, with a maximum width of ~150 Å, and a minimum width of ~70 Å. Scale bar, 100 Å. (G) Unsharpened cryo-EM density of the PSP tau filament displayed as an average of 10 z-slices. Scale bar, 25 Å.

Twisted tau filaments found in PSP superficially appear to have a twofold symmetry (Fig. 1, E and F) but are, in fact, pseudosymmetric ribbons (Fig. 1G). At high-resolution, the compact filament cross-section reveals an asymmetric chain, extending from residues K274-E380 of the tau protein (Fig. 2A, Table S1). The 107 ordered residues adopt a novel fold that includes 11 β-strands (β1-β11, Fig. 2B), with three (β1-β3) from R2 (including K274 from R1), three (β4-β6) from R3, three (β7-β9) from R4 and two (β10-β11) from residues K369-E380. These β-strands are connected by short loops (C291-D295, P301-Q307, C322-G326, K331-G335, P364-K370) punctuated by tight (~75° ± 10°, K281, G304, Q307, C322, G326, P332, G335, G355, G367, K370, H374, L376) or gentle (~20°, S341) turns with breaks in the strand geometry (P312) to relieve strain (Fig. 2, A and B). The residues not involved in β-strand formation facilitate meandering of the chain, similar to other tau filaments (*5–8*).

**Fig. 2.**
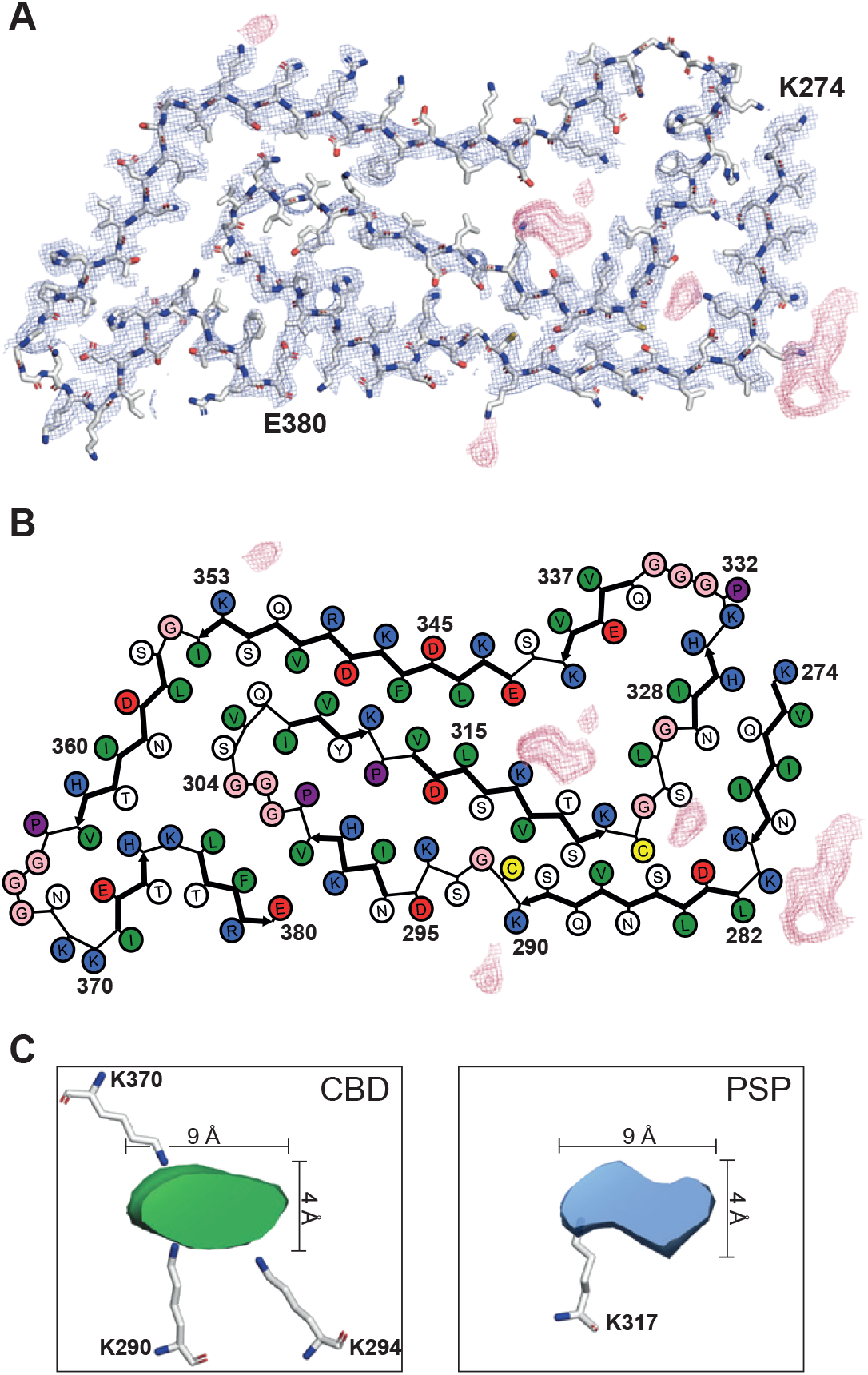
Cryo-EM structure of twisted tau filaments from PSP. A) Cryo-EM density (blue mesh) of PSP tau filament and atomic model (colored sticks). Extra - non-tau protein - densities (pink mesh) directly connected to, or in close proximity of, lysine sidechains are shown. B) Schematic view of the PSP tau filament. Extra densities proximal to K280, K281, K290, K317, and K353 are shown (pink mesh). C) Close-up views and comparison of buried non-tau protein components in CBD (*8*) (green, PDB:6TJO, EMD-10512) and PSP (blue) tau filaments. Despite being bound to distinct lysine residues, both cofactor molecules exhibit very similar dimensions of ~ 9 Å × 4 Å.

The N-terminal heptapeptide K274-K280 (β1) adopts a β-strand conformation similar to tau filaments in CBD (*7, 8*) and forms an antiparallel cross-β interface with N327-H330 (β6) with an inter-sheet separation of ~10 Å (Fig. 2, A and B). There is a tight turn at K281 leading into L282-K290 (β2) which is nestled close to C322 and the tip of β5. Small residues from β2 (S285, V287, S289, C291) are interdigitated with residues C322, S320, and V318 creating an unusually tight interface separated by ~6 Å at the closest points. A compact loop region is formed by C291-D295 preceding β3 (N296-V300) which packs against residues E380 and F378 of β11. The versatile 301PGGG304 forms a straight but compact motif before turning sharply at G304 into a four-residue β-hairpin (304GSVQ307), previously identified in the SufD β-helix (see Methods for details). The short β-strand formed by 308IVYK311 (β4) differs from the other tauopathies as the ubiquitously linear VQIVYK motif (*5–8*) forms part of the right-angled turn. P312 separates β4 and β5 which is formed by residues V313-K321 with a putative salt bridge between D314 and K294. Notably, K317 is attached to a strong, non-tau density that is likely to be a buried cofactor (Fig. 2, A and B). A right-angled turn at C322 introduces a short loop (323GSLG326) to β6 which is formed by 327NIHH330. An extended form of PGGG connects β6 to β7 (Q336-K340). A gentle turn at S341 leads to an extended β-strand formed by E342-I354 (β8) which has closest contact with V309 and K311 of β4. A tight turn at G355 connects to a slightly curved β-strand formed by residues S356-V363 (β9). A loop region adopted by P364-K370 creates a right-angled turn at G367 reminiscent of a similar turn by these seven residues from the tau protein conformer in Pick’s disease (*6*). A short β-strand comprised of residues I371-H374 (β10) packs against both V363-N368 and to T377 *via* an antiparallel cross-β interface with L376-E380 (β11). The extreme C-terminal β-strands (β10 and β11) are connected by K375. Large, non-tau densities decorate the periphery of the filament core and are found in close proximity to K281, K290, and K353 (Fig. 2, A and B). An additional buried density is located close to K280 (Fig. 2, A and B).

Strikingly, a buried cofactor attached to K317 occupies a large cavity in the core of the filament with near-stoichiometric occupancy (Fig. 2, A and B). Unlike the posttranslational modifications which decorate the exposed surface of tau filaments in multiple diseases (*7, 18*), buried non-tau densities may represent small, negatively-charged molecules, e.g., phospholipids, bound to positively-charged lysines. This has been demonstrated for tau filaments from CBD (*7, 8*), and now for PSP (Fig. 2C). The possibility of K317 being posttranslationally modified is unlikely, as mass spectrometry has shown that this site is not ubiquitinated in multiple PSP cases (*18*). The cofactor attached to K317 bears a remarkable similarity in size and shape to a non-tau density connected to K290, K294, and K370 in tau filaments from multiple CBD cases (*7, 8*). Comparison of the densities from CBD and PSP (Fig. 2C) highlights this likeness and points towards the possibility of a common cofactor that potentially helps nucleate and stabilize the core of the filaments in these distinct tauopathies.

Our results show that residues K274-E380 comprise the filament core in two distinct tauopathies - CBD (*7, 8*) and PSP (Fig. 3A). Comparison of the secondary structure motifs adopted by the tau protein in CBD and PSP shows that many contiguous amino acids are identical with on average only one in five residues having a β-strand/loop mismatch (Fig. 3A). This shows that the secondary structure propensities of tau residues K274-E380 are preserved irrespective of subcellular localization, cell-type, or disease etiology (*19, 20*). However, although the secondary structural building blocks are highly similar, the tau conformers in CBD (Fig. 3B) and PSP (Fig. 3C) are radically different both in terms of fold and inter-residue contacts. The molecular-level polymorphism displayed by 4R tau in filaments purified from CBD and PSP human brain is unlike the ultrastructural polymorphism (*21, 22*) found in other tauopathies. In Alzheimer’s disease (AD) (*5*) and chronic traumatic encephalopathy (CTE) (*23*), a disease- and isoform-specific protofilament packs against a second protofilament in one of two ways to form a filament while in Pick’s disease (*6*) and CBD (*7, 8*) it remains single or forms doublet fibrils. Here, we show that the same 4R tau isoforms misfold into two structurally dissimilar protofilaments in CBD (Fig. 3B) and PSP (Fig. 3C) and find no evidence of the twisted tau filament in PSP laterally associating with another (i.e., no ultrastructural polymorphism).

**Fig. 3.**
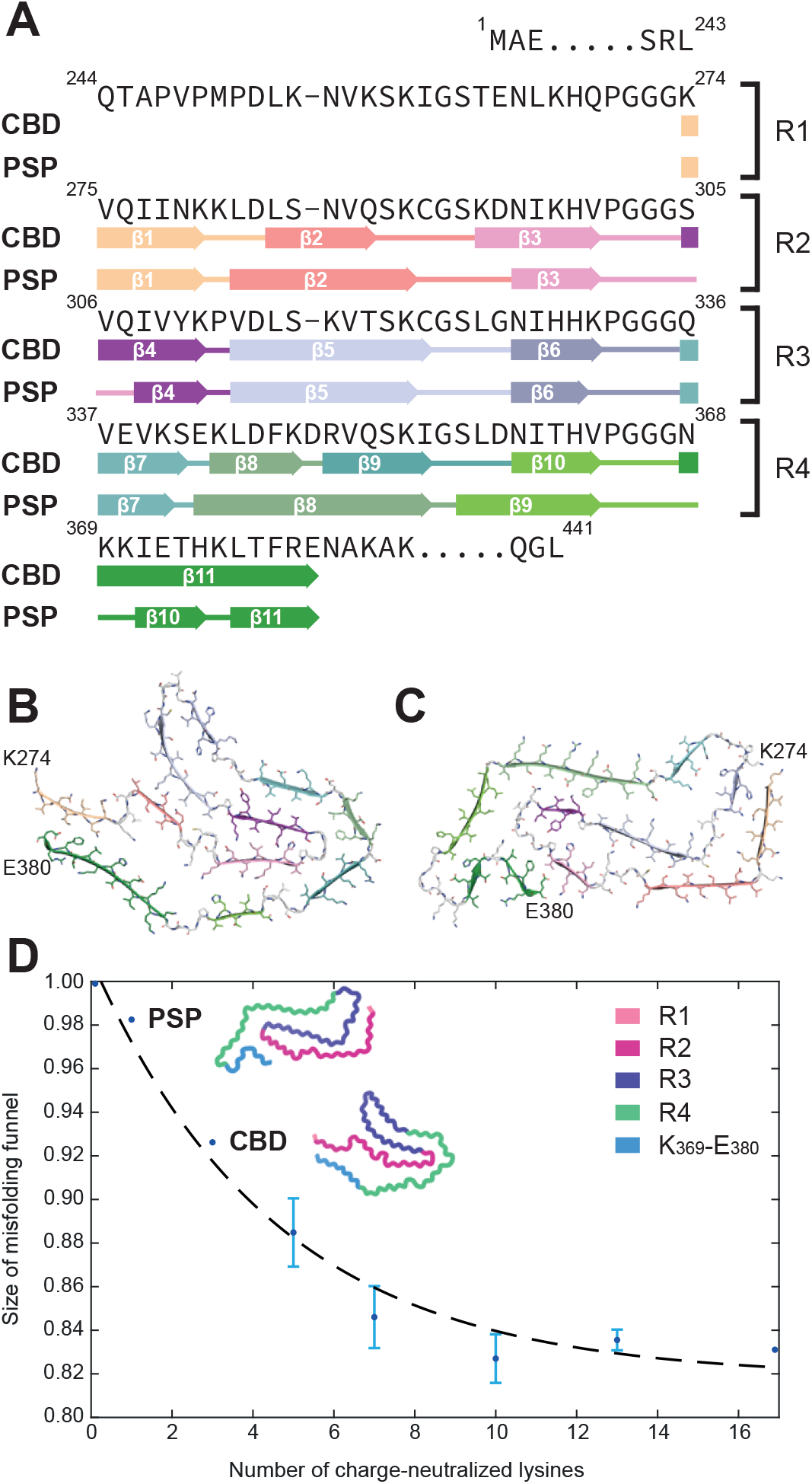
Structural comparison between CBD and PSP tau filaments and predicted effect of charge-neutralized lysines on tau misfolding. A) Comparison of tau filament core sequence and structural motifs in CBD and PSP. The sequence is organized into four microtubule-binding repeats (R1-R4) and K369-E380, with the positions of β-strand and loop regions shown. B) and C) Atomic models of CBD singlet (B, PDB: 6VHA) and PSP (C) filaments, with β-strands colored according to (A). D) Plot of tau misfolding funnel size as a function of the number of charge-neutralized lysine sidechains. Fractional change in the width (s.d.) of the free energy minimum is a slowly decaying function of the number of lysines that are rendered more hydrophobic *via* charge neutralization by cofactor binding or posttranslational modification. The predicted effect of cofactor binding to K317, or to K290, K294, and K370, in tau filaments from PSP and CBD, respectively, is shown. The remaining points on the curve represent random patterns of posttranslational modification or cofactor binding to any of the 17 lysine sidechains in K274-E380. For reference, tau filaments in CBD have a total number of nine charge-neutralized lysines arising from cofactor binding and posttranslational modification (*7*). Schematics of the tau filament structures from CBD and PSP are shown to scale.

Considering that the structural building blocks are so similar (Fig. 3A), why are the misfolded structures of 4R tau in PSP and CBD so different (Fig. 3, B and C)? Given that many surface-exposed lysines in tau filaments from CBD are buried in the core of PSP (K317, K321, K340, K375, Fig. 2, A and B; Fig. 3C) and vice versa (K281, K290, K298, K370, Fig. 3B), different patterns of posttranslational modifications (*18*) may influence the tau filament structure (*7*). The location and charge of buried co-factors could also play a significant role in nucleating and perpetuating a specific tau fold, particularly since the buried co-factor in tau filaments is bound to different residues in PSP (K317) and CBD (K290, K294, and K370) (*7, 8*) (Fig. 2C). Acetylation, ubiquitination, or cofactor binding will neutralize the positive charge of lysine sidechains in the tau filament core; to explore the effect of these perturbations, we simulated tau misfolding using a simple folding energetics model (see Methods for details).

Since the structures of individual tau molecules in filaments from PSP and CBD are largely planar, we treat this as a simplified 2D folding problem (*24, 25*). The planarity of the tau conformers (Fig. 3, B and C) drastically reduces the number of degrees of freedom while the secondary structure propensities of the tau molecule (Fig. 3A) limit its conformational space even further. Using the simulated folding of K274-E380 as a reference, we normalize the size of the misfolding funnel to unity and find that as we increase the number of charge-neutralized lysines in K274-E380 the radius of gyration of the tau molecule remains unchanged but the width of the misfolding funnel decreases to 0.8 (Fig. 3D). This narrowing of the free energy minimum by up to 20% reflects the reduction in conformational space upon binding of cofactors to, or posttranslational modifications of, lysine residues in the misfolded tau molecule. We propose that cofactor binding and posttranslational modifications to the tau molecule alter the free energy landscape of tau misfolding, increasing the likelihood of a small set of energetically favorable structures as detected in human disease (*5–8, 23*).

The molecular polymorphism displayed by 4R tau explains why many positron emission tomography (PET) ligands do not specifically bind to tau filaments from PSP with high affinity (*26*). The structure shown here (Fig. 2, A and B; Fig. 3C) identifies many potential binding sites for small ligands that could be targeted *via* structure-based design (*27*). Significantly, R3 is buried in the core of the filament and is not accessible (Fig. 2, A and B; Fig. 3C). Approximately half of the amino acid sidechains in R2 and R4 are exposed on the surface of the filament (Fig. 2, A and B; Fig. 3C). Since PSP tau filaments have fewer posttranslational modifications than other tauopathies and are more prone to enzymatic digestion (*18*), these surface-exposed amino acid sidechains are primed for the development of diagnostic and therapeutic small molecules.

The structures of tau conformers from many neurodegenerative diseases including AD (*5*), Pick’s disease (*6*), CTE (*23*), and CBD (*7, 8*) have shown that each tauopathy has a unique structural signature. What has emerged is that the tau protein aggregates in an isoform-specific manner, adopting unique shapes that define the templated growth of filaments in 3R, 3R/4R, and 4R tauopathies (*28*). Mediating the structural diversity of these tau filaments are disease-specific patterns of posttranslational modifications (*7, 18*) and, in at least three tauopathies (CTE (*23*), CBD (*7, 8*), and PSP), buried cofactor molecules. This study has identified a novel form of structural diversity – molecular polymorphism – whereby the same 4R tau isoforms adopt two dramatically different folds. It could be argued that tau protofilaments from AD (*5*) and CTE (*23*) also display molecular polymorphism, since these protofilaments both contain 3R/4R tau isoforms; their folds, however, are highly similar (*23*). Our work demonstrates that tau, while not a metamorphic protein (*29*) (since molecular recycling occurs over many days (*30*)) adopts two vastly dissimilar structures within diverse cellular environments manifesting in phenotypically distinct neurodegenerative diseases.

## Acknowledgments

The authors are grateful to Dr. R. S. Mann for critical reading of the manuscript, and to the patients and their families for their participation in this work. Human biological samples and associated data were obtained from the Mayo Clinic Brain Bank. We are grateful to Dr. X. Chen and Dr. K. Song for help collecting data on the Titan Krios at the University of Massachusetts Medical School Cryo-EM Core Facility; R. Grassucci and Dr. Z. Zhang for help collecting data at the Columbia University Cryo-Electron Microscopy Center including the Titan Krios housed at the Zuckerman Institute and the Titan Krios housed at the Simons Electron Microscopy Center and National Resource for Automated Molecular Microscopy (New York Structural Biology Center). This paper is dedicated to the memory of Mr. Patrick Trainor.

## Author contributions

A.W.P.F. conceived the experiments. D.W.D. and M.D. identified patients and performed neuropathology. Y.C. purified tau filaments from PSP cases. A.W.P.F. performed cryo-EM experiments. A.W.P.F. and T.A. processed cryo-EM data. A.W.P.F. and X.X. built atomic models. A.W.P.F. and S.D. performed and analyzed simulations. A.W.P.F., X.X., T.A., S.D., M.T., C.C., D.W.D., and L.P. prepared the manuscript. A.W.P.F. supervised the project.

## Data and materials availability

Density maps and coordinates for the twisted tau filament from PSP human brain will be deposited in the Electron Microscopy Data Bank (EMDB) with ID code EMDB-XXXXX and the Protein Data Bank (PDB) with ID code XXXX, respectively, upon acceptance of this manuscript for publication.

## Materials and Methods

### Clinical history and neuropathology of PSP case

The patient was a 68-year-old white man, an electrical engineer by profession, with no significant past medical history when he developed neurological deterioration six years prior to death. Initial symptoms were double vision. He subsequently developed slurred speech and bradykinesia. He had difficulty walking and unexplained falls. He was diagnosed with Parkinson-plus syndrome three years before he died. He was treated with dopamine replacement therapy without any significant benefit. On examination, he had no significant cognitive impairment, but he had postural instability and rigidity most marked in the right arm and hand. Corticobasal degeneration was in the clinical differential diagnosis. In the last year of life his markedly asymmetric neurologic deficits suggested the possibility of a stroke. Post-mortem neuropathologic evaluation of the left half of the brain revealed findings of cortical-predominant PSP (*31*), with severe neuronal and glial tau pathology in the motor and premotor cortices, and marked white matter oligodendroglial pathology at the gray-white junction. There was also primary age-related tauopathy, with sparse medial temporal Alzheimer type neurofibrillary tangles, but no cerebrovascular pathology. The brain weight was 1,340 grams. Postmortem genotyping revealed him to be APOE E2E3 and MAPT H1H1.

### Immunohistochemistry

Formalin-fixed brains undergo systematic and standardized sampling (*32*), including at least sixteen sections from the formalin-fixed hemibrain. Tissue samples are processed for paraffin embedding in a TissueTek VIP 6 (Sakura). Paraffin-embedded tissue sections are cut at a thickness of 5 μm and mounted on glass slides. The glass-mounted tissue sections are deparaffinized in xylene and then hydrated in decreasing concentrations of ethanol. The deparaffinized tissue sections are pretreated with steam in deionized water for 30 minutes. Immunohistochemistry for tau utilizes a mouse monoclonal antibody, CP13 (IgG2b, phospho-serine 202 www.alzforum.org/antibodies/tau-phos-ser202-cp13), at 1:1,000 dilution. CP13 was provided by the late Dr. Peter Davies, Feinstein Institute, North Shore Hospital, NY. Immunohistochemistry is performed on a DAKO Autostainer with DAKO EnVision reagents (DAKO, Carpinteria, CA). Immunostained slides are counterstained with hematoxylin and coverslipped.

### Purification of tau filaments from PSP human brain

To obtain a sufficient yield of tau filaments from PSP tissue, we evaluated the amount of tau present within the sarkosyl-insoluble (P3) fraction from four different brain regions in three different PSP cases, without pronase-treatment to preserve posttranslational modifications (*7, 33*). As tau levels were consistently highest in the P3 fractions from caudate and frontal cortex, we screened these two regions in an additional 12 PSP cases. Using EM, we observed a good yield of filaments (~0.006 mg/mL) with low background in the P3 fraction from the caudate of case #1. To further characterize the PSP case used for cryo-EM, we evaluated total and phosphorylated tau levels in the P3 fraction of case #1 and 3 additional PSP cases by western blot (Fig. 1D). The similarities in positivity for common disease-associated phosphoepitopes indicates case #1 is a typical PSP case, suggesting that the morphology of tau filaments in this case will be representative of PSP.

### Cryo-electron microscopy of tau filaments from PSP human brain

3 μL aliquots of purified tau filaments from PSP human brain tissue (~0.006 mg/mL) were applied to glow-discharged holey carbon grids (Quantifoil Au R1.2/1.3, 300 mesh), blotted with filter paper and plunge-frozen into liquid ethane using an FEI Vitrobot Mark IV. Images of the frozen grids were collected on three FEI Titan Krios microscopes on multiple occasions due to the paucity of the tau filaments. Imaging conditions were always identical (Table S1), but minute differences in Å/pixel had to be corrected for during image processing. All images were acquired at 300 kV and energy filtered (slit width of 20 eV) using a Gatan K2/K3 Summit detector in counting mode. All movies were collected at a nominal pixel size of 1.0605 Å, using a dose rate of 1.5 electrons per Å^2^ per frame and a defocus range of −1 to −3 μm (Table S1).

### Helical reconstruction

Preprocessing of movie frames was performed using MotionCor2 (*34*) and Gctf (*35*). Helical processing of tau filaments from PSP was mainly performed using RELION (*36*), as previously described (*5*). Due to an uneven distribution of views (fig. S1A), helical reconstruction of the final particle stack was non-uniform refined in cryoSPARC (*37*) version 3.0.0. Twisted tau filaments were manually selected and segments were extracted at various Å/pixel (1×, 2×, 3×, 5× binned) and 2D classified to determine the helical parameters and ascertain whether - or not - the filament had C2 symmetry. Initial models were generated from 2D class averages of helical pitch views and used as a reference for helical reconstruction in C1 using 3D classification with a T regularization value of 100. When a clear chain trace emerged in the helical cross-section at 1.0605 Å/pixel, a final subset of particles was 3D auto-refined with T = 20. Helical twist and rise were optimized (Table S1) with the tilt angles showing a unimodal distribution around 90°, as expected. Once the helical parameters were optimized, helical symmetry remained fixed in all subsequent refinements or 3D classifications. Masking and particle subtraction was used to auto-refine and 3D classify sub-volumes of the filament to improve density in specific regions (e.g. K274-K331 and Q307-V363). Once the helical parameters were established, particle stacks were also refined in cryoSPARC version 3.0.0 (*37*) applying helical symmetry and using non-uniform refinement. In addition to helping with the uneven distribution of views, helical reconstruction in cryoSPARC version 3.0.0 (*37*) ensured that there was not overfitting of the data. Postprocessing was performed in RELION (*38*), cryoSPARC version 3.0.0 (*37*), and DeepEMHancer (*39*) *via* COSMIC (*40*) also aiding interpretation of the maps. Local resolution estimates were determined using RELION (*38*) and displayed using Chimera (*41*) (fig. S1B). The final overall resolution of 3.9 Å was calculated from Fourier shell correlations at 0.143 between the two independently refined half-maps (*42*) in cryoSPARC version 3.0.0 (*37*) to ensure against overfitting (fig. S1C).

### Model building and refinement

First, a continuous 107 amino acid polyalanine chain was constructed into one rung of the sharpened density, with backbone geometries refined using real-space refinement in COOT (*43*). Various cryo-EM maps were used, or combined, for model-building including maps generated by RELION (*36*), cryoSPARC version 3.0.0 (*37*), and postprocessed in RELION (*38*), cryoSPARC version 3.0.0 (*37*), or DeepEMhancer (*39*). Subtracted maps of regions K274-K331 and Q307-V363 were also used to define the Cα positions. Landmark amino acid side chains (Y310, F346, R349, Q351, K353, H362), regions with contiguous bulky sidechains (N296-P301, L376-E380), and an unusually tight interdigitation of seven residues (S285, V287, S289, C291 with residues C322, S320, V318) enabled the discrimination between each of the tandem repeats (R1-R4) and K369-E380. Matching segments were then taken directly from published atomic models and rigid body fitted into the sharpened density, followed by real-space refinement in Phenix 1.19.1-4122 (*44*) using secondary structure and hydrogen bond restraints of the β-sheets. Briefly, K274-K280 was taken from PDB-entry 6TJO (*8*); K281-V283, C291-P301, Q307-C322, G326-K331, Q336-K340, and L376-E380 were taken from PDB-entry 6VHA (*7*); G323-L325 was taken from PDB-entry 5O3L (*5*); L344-V363 and I371-H374 were taken from PDB-entry 6GX5 (*6*). Some segments were docked and mutated, and the 107-residue chain was assembled into a nine-chain stack in Chimera (41) followed by real-space refinement in Phenix 1.19.1-4122 (*44*). Segment D348-I354 in PDB-entry 6GX5 (*6*) was mutated to L284-K290 in PSP. The four-residue β-hairpin formed by 304GSVQ307 was modelled from E289-C295 in the SufD β-helix (1VH4) (*45*). A few residues overlapped with previously docked β-strands and retained for alignment purposes only to be removed later to form a continuous chain. The extended and curved conformation of 332PGGG335 and 364PGGGNKK370 were modelled using motifs from PDB-entry 6GX5 (*6*). Clashes were resolved in COOT (*43*) and ISOLDE 1.0b3.dev7 (*46*) followed by real-space refinement in Phenix 1.19.1-4122 (*44*) using a resolution limit of 4.0 Å. All structural figures were generated using PyMOL (*47*), Chimera (*41*) or ChimeraX (*48*), and atom2svg.py (*49*).

### 2D tau protein folding simulations

For two-dimensional protein folding simulations we used an Ant Colony Optimization (ACO) algorithm kindly provided by Drs. A. Shmygelska and H. H. Hoos (*25, 50*) which predicts a protein’s conformation from its amino acid sequence. It has been shown previously that the Hydrophobic-Polar (HP) model (*24*) can approximately capture β-sheet folding energetics (*51*). The amino acid sequence K274-E380 was first converted into a HP model (*24*) and subsequently the ACO algorithm was applied to it. The ACO algorithm outputs multiple possible folding configurations of the sequence sorted by their free energy. Each of these folding configurations was mapped onto an *x-y* plane and their radius of gyration was calculated. This process was iterated multiple times, and the radius of gyration was plotted against the free energy of each configuration. Lastly, the mimima were fitted using a 2D Gaussian function to find the standard deviation of the free energy basins. We conducted this process on our 107 amino acid sequence which contains 17 polar lysines. To simulate the binding of negatively-charged cofactors to, or posttranslational modifications of, lysine sidechains, we substituted specific lysine residues with non-polar amino acids (since charge-neutralization of lysine increases its hydrophobicity) and observed the changes in the protein folding model. As expected, in the wild-type structure, which has all lysines intact, the standard deviation of the energy minimum was the highest, indicating many possible configurations. For the PSP structure we substituted one cofactor-binding lysine (K317) which decreased the standard deviation of the free energy. Following the trend, for CBD we substituted 3 cofactor-binding lysines (K290, K294, and K370) and obtained an even smaller standard deviation. The standard deviation continued to decrease for 5, 7, and 10 randomly substituted lysine residues (10 random permutations per data point to generate error bars), eventually plateauing between 10 and 17 residues. Our simulations showed that as more lysines were substituted for hydrophobic residues the variability in free energy minima decreased. This trend indicates that having cofactors bound to the molecule, which we indicated by more substitutions of polar lysines with non-polar amino acids, allows fewer folding possibilities.

**Table S1:** Cryo-EM data collection, model reconstruction, refinement and validation statistics.

| Cryo-EM data collection, reconstruction, refinement and validation statistics |  |
| --- | --- |
| **Data Collection** |  |
| Microscope | FEI Titan Krios G3 |
| Camera | Gatan K2/K3-Summit |
| Voltage (kV) | 300 |
| Magnification | ×105,000/×81,000 |
| # of frames (no.) | 40 |
| Total electron dose (e<sup>-</sup>/Å<sup>2</sup>) | 60 |
| Defocus range (μm) | -1 to -3 |
| Pixel size (Å) | 1.0605 |
| **Reconstruction** |  |
| Box size (pixel) | 270 |
| Final particle images (no.) | 1,966 |
| Segments extracted (no.) | 18,709 |
| Estimated accuracy of translations (Å, FSC=0.143) | 0.77 |
| Estimated accuracy of rotations (Å, FSC=0.143) | 1.41 |
| Symmetry imposed | C1 |
| Map resolution (Å, FSC=0.143) | 3.9 |
| Map sharpening B-factor (Å<sup>2</sup>) | -142 |
| Helical twist (°) | -0.75 |
| Helical rise (Å) | 4.73 |
| **Refinement and Validation** |  |
| Model composition (1 chain) |  |
| Non-hydrogen atoms | 813 |
| Protein residues | 107 |
| R.m.s deviations (stack of 9 chains) |  |
| Bond lengths (Å) | 0.003 |
| Bond angles (°) | 0.858 |
| Validation (MOLPROBITY) |  |
| Clashscore (stack of 9 chains) | 16.2 |
| Poor rotamers (%) | 0 |
| Cβ deviations (%) | 0 |
| Ramachandran plot (stack of 9 chains) |  |
| Favored (%) | 90 |
| Allowed (%) | 9.1 |
| Disallowed (%) | 0.9 |

**fig. S1.**
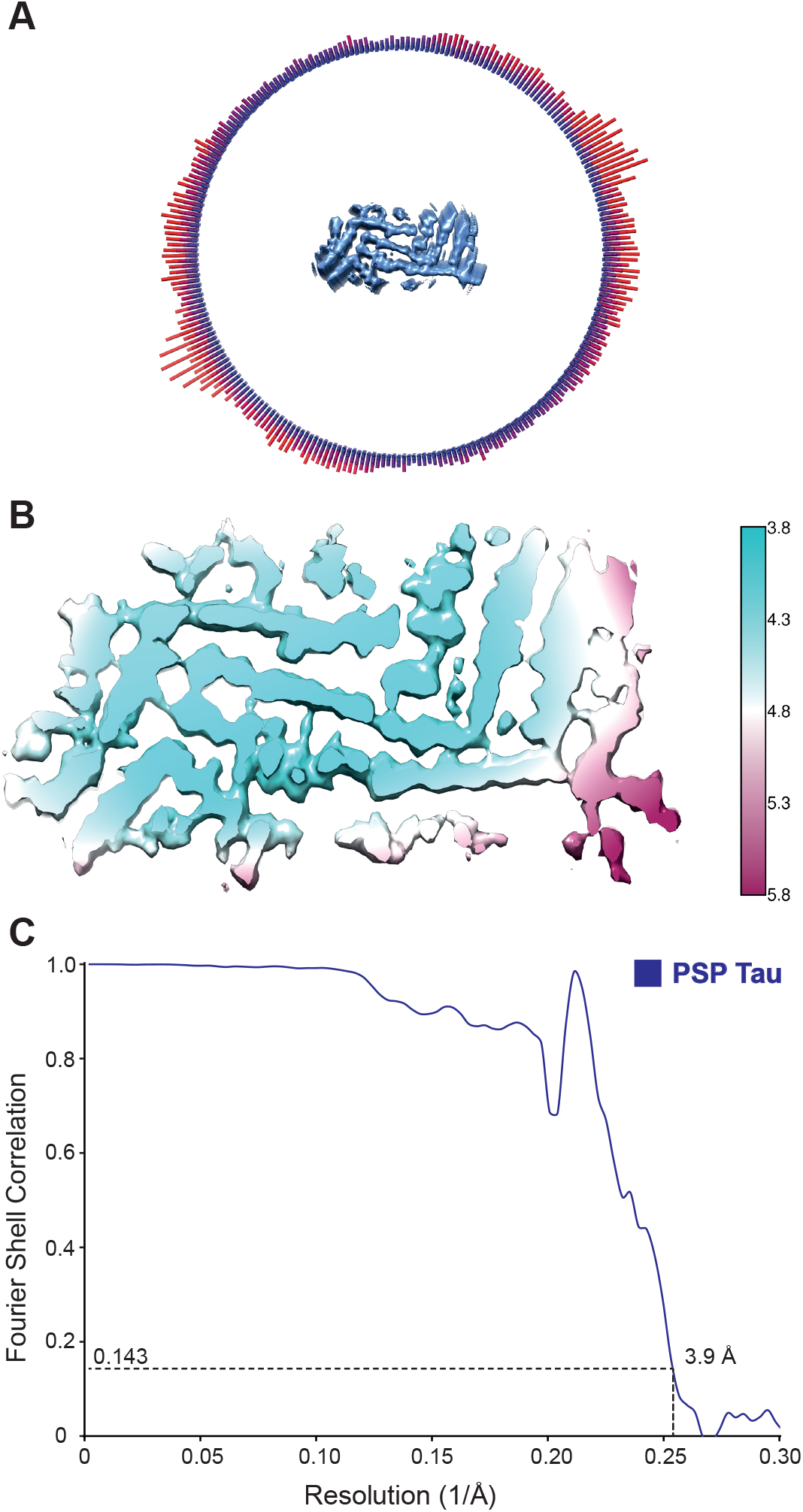
Resolution estimates and cryo-EM validation of the PSP tau filament map. A) Angular distribution of number of filament particles for a given viewing direction (from few to many, blue to red) with respect to the unsharpened helical cross-section (blue surface). B) Local resolution estimates of the PSP tau filament map ranging from cyan (3.8 Å) to maroon (5.8 Å). C) Independently refined half-maps Fourier shell correlation (FSC) curve (blue solid line).

